# Cyano-assassins: Widespread cyanogenic production from cyanobacteria

**DOI:** 10.1101/2020.01.04.894782

**Authors:** Manthos Panou, Spyros Gkelis

## Abstract

Cyanobacteria have been linked with hydrogen cyanide, based on their ability to catabolize it by the nitrogenase enzyme, as a part of nitrogen fixation. Nitrogenase can also use hydrogen cyanide instead of its normal substrate, dinitrogen and convert it to methane and ammonia. In this study, we tested whether cyanobacteria are able, not only to reduce, but also to produce HCN. The production of HCN was examined in 78 cyanobacteria strains from all five principal sections of cyanobacteria, both non-heterocytous and heterocytous, representing a variety of lifestyles and habitats. Twenty-eight (28) strains were found positive for HCN production, with universal representation amongst 22 cyanobacterial planktic and epilithic genera inhabiting freshwater, brackish, marine (including sponges), and terrestrial (including anchialine) habitats. The HCN production could be linked with nitrogen fixation, as all of HCN producing strains are considered capable of fixing nitrogen. Epilithic lifestyle, where cyanobacteria are more vulnerable to a number of grazers and accumulate more glycine, had the largest percentage (75%) of HCN-producing cyanobacteria compared to strains from aquatic ecosystems. Further, we demonstrate the isolation and characterisation of taxa like *Geitleria calcarea* and *Kovacikia muscicola*, for which no strain existed and *Chlorogloea* sp. TAU-MAC 0618 which is, to the best of our knowledge, the first bacterium isolate from anchialine ecosystems. Our results highlight the complexity of cyanobacteria secondary metabolism, as well as the diversity of cyanobacteria in underexplored habitats, providing a missing study material for this type of environments.

## 1. Introduction

Cyanobacteria are ubiquitous photosynthetic prokaryotes, found literally in any illuminated environment and unexpectedly in some dark subsurface ones (Hubalek et al., 2016; Puente-Sánchez et al., 2018). This phylum predominated Earth, when the environment was still reductive ca. 1.3 billion years prior to the great oxidation event (Gumsley et al., 2017). Cyanobacteria often produce secondary metabolites, in response to biotic or abiotic stress in the surrounding environment, providing protection and aiding in survival over other species (Gupta et al., 2013; Singh et al., 2016).

Cyanobacteria are considered to be the ancestors of chloroplasts genome (Allen, 2015). For example, plants have probably obtained gene copies implicated in a variety of biosynthetic pathways through early horizontal gene transfer from proteobacteria and cyanobacteria (Timmis et al., 2004). A number of secondary metabolites (e.g. many terpenoids, quinolizidine alkaloids, piperidine alkaloid coniine) are produced completely or partly in chloroplasts and/or mitochondria (Wink, 2003, 2008). Moreover, there is an indication that many enzymes and pathways are common in plant and cyanobacteria secondary metabolite production (Chen et al., 2016; Nielsen et al., 2016).

The term “cyanide” is used loosely and refers to both the cyanide anion (CN-) and undissociated hydrogen cyanide (HCN) (Knowles and Bunch, 1986). Cyanides are present in various environmental elements such as water, soil, food and biological materials like blood urine and saliva at the levels of micrograms per litre to milligrams per litre (Dzombak et al., 2006; Barceloux, 2009). Considering the presence of cyanide in various parts of the inanimate environment and biota as well as their toxicity, there is no doubt on increasing demand for information on their prevalence in the elements of the environment (Dzombak et al., 2006). This compound has been found in some foods and seeds in the amounts above the limit recommended by WHO and FAO (Gernah et al., 2011). The pKa of cyanide is 9.3; it is therefore present largely as HCN at neutral pH values (Eisler and Wiemeyer, 2004). HCN is volatile (boiling point 26°C) and is less dense than air. Hence, cyanide formed by microbial cultures will be rapidly lost to the environment. Cyanide is largely toxic for aerobic cell metabolism as it binds to the mitochondrial cytochrome oxidase a3 enzyme; binding of cyanide to the ferric ion of the cytochrome oxidase a3 inhibits the terminal enzyme in the respiratory chain and halts the electron transport and oxidative phosphorylation, which subsequently leads to intracellular hypoxia (Hall, 2007).

Despite its toxicity, cyanide is a natural compound synthesized by a variety of organisms, including bacteria, fungi, plants, and animals, in which cyanogenesis may serve as defensive or offensive mechanism (Luque-Almagro et al., 2016). The HCN synthase required for bacterial cyanogenesis is expressed during transition from exponential to stationary phase of growth under oxygen limitation in response to the FNR-like anaerobic regulator ANR (Luque-Almagro et al., 2016). On the other hand, many microorganisms have evolved enzymatic pathways for cyanide degradation, transformation, or tolerance, and many of them are even able to use cyanide as a nitrogen source for growth (Luque-Almagro et al., 2016; Kumar et al., 2017; Park et al., 2017).

To date the green alga *Chlorella vulgaris* and the cyanobacteria *Anacystis nidulans* (=*Asterocapsa nidulans*), *Plectonema borganum*, and *Nostoc muscorum*, are the only photosynthetic micro-organisms known to be cyanogenic (Pistorius et al., 1979; Vennesland, 1981). However, it is likely, given the nature of the cyanogenic pathways involved, that many more of these micro-organisms to be cyanide producers. Photosynthetic micro-organisms synthesize cyanide from a wide range of metabolites by at least two distinct systems (Vennesland et al., 1982) (i) the amino acid oxidase-peroxidase system and (ii) the glyoxylic oxime system.

Diazotrophic cyanobacteria have been linked with hydrogen cyanide, based on their ability to catabolize it by nitrogenase, an enzyme normally responsible for the reduction of N_2_. Nitrogenase can also use hydrogen cyanide instead of its normal substrate, dinitrogen and convert it to methane and ammonia (Gantzer and Maier, 1990). The N_2_-fixing cyanobacterium, *Anabaena* is able to biodegrade cyanides and produce CH_4_ in batch reactors. Gantzer and Maier (Gantzer and Maier, 1990) showed that *Anabaena* reduced cyanides by nitrogenase to CH_4_ and NH_3_. The rate for CH_4_ production was ten times faster than expected based on literature. However, in these cases, the assumption was made based on induced HCN in batch reactor.

In this study, we tested whether cyanobacteria are able, not only to reduce, but also to produce HCN, broadening our knowledge on the biosynthesis of this unique molecule. We examined HCN production in representative genera from all five of the principal sections of cyanobacteria, in both non-heterocytous and heterocytous species, representing a variety of lifestyles and habitats. We also correlate the production of HCN with epilithilic lifestyle as the main hypothesis for the HCN production from cyanobacteria.

## 2. Materials and Methods

### 2.1 Cyanobacterial Strains

Seventy-eight (78) strains of cyanobacteria, representing five cyanobacteria orders, with different morphological features and a wide variety of habitats and lifestyles, were used in this study (Table S1). Twenty-nine strains were isolated from different freshwaters of Greece (Gkelis et al., 2019), nine of them were isolated from sponges (Konstantinou et al., 2018), whereas the rest 40 strains were isolated in this study from various environments (terrestrial caves, coastal areas, thermal springs, brackish and freshwater systems) across Europe, between 2013 and 2018. Strains were isolated on solid growth media using classical microbiological techniques (see (Gkelis et al., 2005, 2015)), purified by successive transfers and using antibiotics (such as cycloheximide and ampicillin) as described in Rippka (Rippka, 1988); all strains were derived from a single colony or trichome. The cultures were grown as batch clonal unialgal cultures in BG11, with or without (for the nitrogen-fixing strains) nitrogen, and MN medium (Rippka, 1988). All strains are deposited in the Thessaloniki Aristotle University Microalgae and Cyanobacteria (TAU-MAC) culture collection (Gkelis and Panou, 2016) and maintained at 20 ±1°C or 25 ±1°C (for strains of the genera *Desertifilum* and *Calothrix*) with a light intensity of 25μmol m^−2^s^−1^ and with a light/dark cycle of 12:12h.

### 2.2 Cyanogenesis analysis

In order to assess the ability to produce HCN and estimate the relative frequency of cyanogenesis (proportion of cyanogenic versus acyanogenic cyanobacteria strains), we screened each strain for the presence of HCN using the Feigl-Anger assay, which determines the presence or absence of HCN in a sample using a colour change reaction (Gleadow and Møller, 2014). In this assay, the dried paper turns blue following oxidation of the tetra base when it is exposed to the HCN gas that is produced. For the extraction of HCN, we modified the original protocol (Thompson et al., 2016), implemented in plants and includes freeze-thaw, as freeze-thaw is not an efficient method for cell lysis in cyanobacteria (Kim et al., 2009). Cyanobacteria cells were harvested at the exponential growth phase (between 30-45 days of growth) by centrifugation of 1.5 mL culture material. The supernatant was removed and the pellet was dissolved in 0.8 mL of Lysis Buffer (2% w/v Cetyl trimethylammonium bromide, 100 mM Tris-HCl, 1.4 M NaCl, 1% w/v Polyvinylpyrrolidone, 20 mM, Na_2_EDTA 0.2% w/v LiCl, pH 8). The same procedure was repeated with the addition of dH_2_O instead of Lysis Buffer. A 96-well plate was filled with 80 μL of cyanobacteria isolates extract to each well. We secured Feigl-Anger test paper over the plates, incubated the plates for 1.5 h at 37 °C, and then scored each well for cyanide, which is indicated by a blue colour (Figure S1). A permanent record of the detection paper was made by scanning the paper immediately after exposure because the blue color fades with time. As positive control, 4 cm^2^ of leaf tissue of a cyanogenic individual of the plant *Trifolium repens* (Deligiannis et al., 2018) was used.

### 2.3 Nitrogen fixation capability

Cyanobacteria strains were also evaluated for the ability to perform nitrogen fixation by PCR targeting one gene fragment (*nifH*) of the nitrogenase gene cluster (Table S2). The reactions were performed according to Panou et al. (Panou et al., 2018). Thermal cycling was carried out using an Eppendorf MasterCycler Pro (Eppendorf). PCR products were separated by 1.5% (w/v) agarose gel in 1X TAE buffer. The gels were stained with Midori Green Advanced (NIPPON Genetics Europe GmbH) and photographed under UV transillumination.

### 2.4 Polyphasic taxonomy

The cyanobacteria isolates, which were positive for the production of HCN, were characterised based on their morphology and their phylogeny as described in Gkelis et al. (Gkelis et al., 2019). Briefly, strains were identified based on their morphology using the taxonomy books by Komárek and Anagnostidis (Anagnostidis and Komarek, 1988; Komarek and Anagnostidis, 1989; Komárek and Anagnostidis, 2008) and Komárek (Komárek, 2013). The phycocyanin operon and the internal genetic spacer (*cpcBA*-IGS), 16S rRNA gene, and the 16S-23S rRNA internal transcribed spacer (ITS) were used to assess the molecular phylogeny of the strains. PCR was carried out on using the primer pairs shown in Table S2 and PCR conditions described in detail in Gkelis et al. (Gkelis et al., 2019). Sequence data were obtained by capillary electrophoresis (GENEWIZ, Takeley, UK). The obtained nucleotide sequences were edited with Unipro UGENE 1.29.0 (Okonechnikov et al., 2012). Nucleotide sequences were deposited in GenBank database of the National Center for Biotechnology Information (NCBI) (Table S3). Sequences were blasted and the closest relative(s) for each sequence were included in the phylogenetic trees. For the phylogenetic analyses, we selected sequences (>1200 and >600 bp, for 16S-23S rRNA and *cpcBA*-IGS, respectively) in order to examine phylogenetic position of our strains. The phylogenetic analyses were conducted with Mega (V7.0) software (Kumar et al., 2016). Multiple sequence alignments were conducted using the CLUSTALW software. All missing data and gaps were excluded from the analysis by choosing the complete deletion option. A consensus phylogenetic tree were constructed using maximum likelihood (ML). The best fitting evolutionary models for the ML analyses were the Tamura 3-parameter + G model for all the targets analysed. Bootstrap replicates (n=1,000) were performed. Phylogeny was also inferred with Bayesian Inference (BI) phylogenetic approach with MrBayes (V3.2.6) software (Ronquist et al., 2012). The general time-reversible (GTR) with gamma distribution of rates and a proportion of invariable sites evolutionary model was selected by applying PAUP* (V5.0) (Swofford, 2002). Bayesian analysis consisted of two independent Markov Chain Monte Carlo runs, performed by four differentially heated chains of 10 × 10^6^ generations and trees were sampled from the chain every 1000 generations. All phylogenetic trees were visualized via the FigTree (V1.4.3) software (Rambaut).

## 3. Results

### 3.1 HCN Production

Twenty-eight (28) cyanobacteria strains (Table 1) were found positive for HCN production (Figure S1). Cyanobacteria strains, with different lifestyles, were found capable of producing HCN (Figure 1). Strains, isolated from mats in rocks, were found to be positive in HCN production with a relative frequency of 75 %. Three strains (50 % percentage) from brackish environments were found positive for the production of HCN, whilst cyanide was found to be present also in sponge-associated cyanobacteria, even though in low percentage (only 22% of the sponge symbiotic cyanobacteria tested). The habitat, with the lowest number of HCN-producing isolates, was freshwater ecosystems (Table 1, Figure 1), where only the 13 % of freshwater cyanobacteria were positive for HCN production.

**Table 1.**
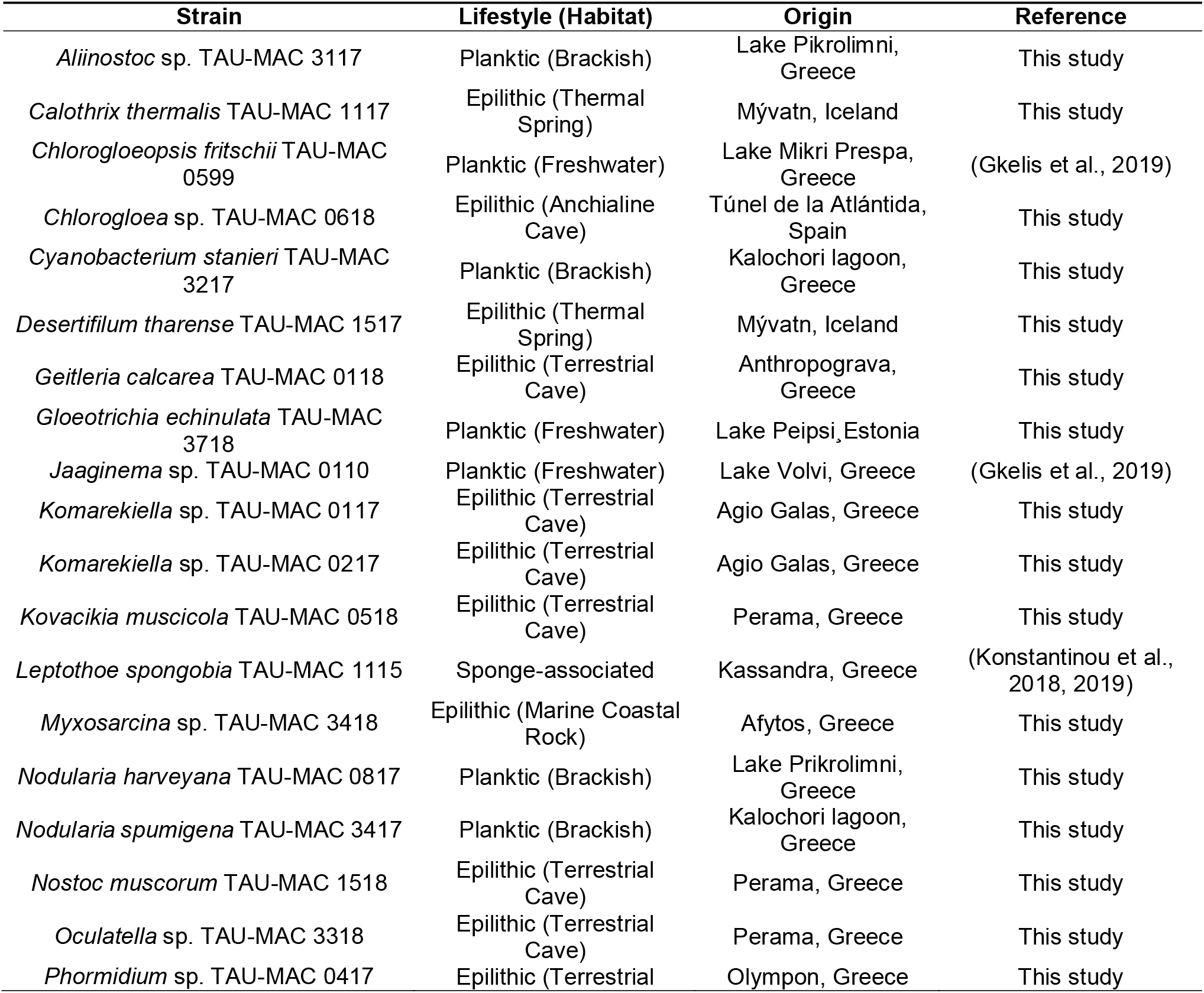
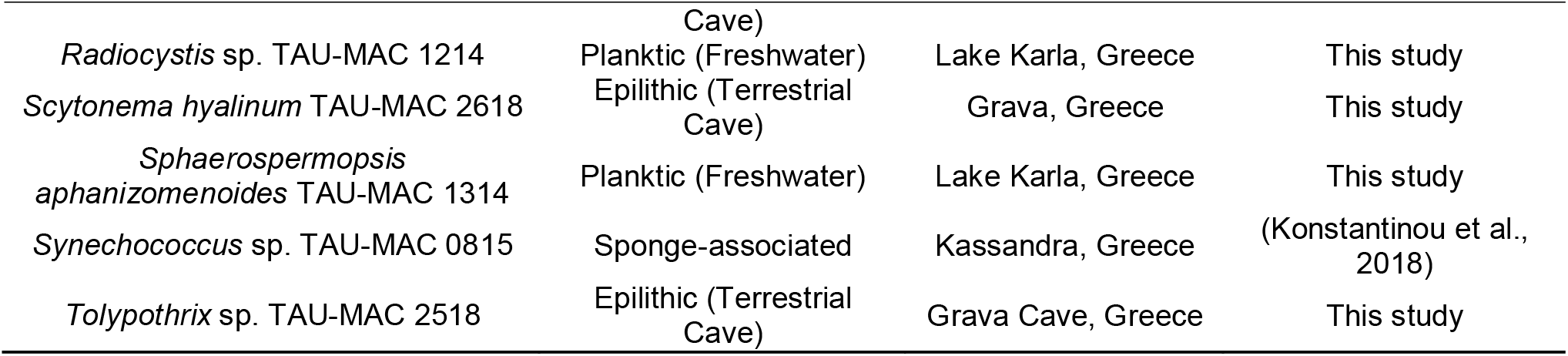
Cyanobacteria strains of TAU-MAC collection positive for HCN production, their origin, habitat, and lifestyle. Table S3 contains the complete list of strains tested for HCN production.

**Figure 1.**
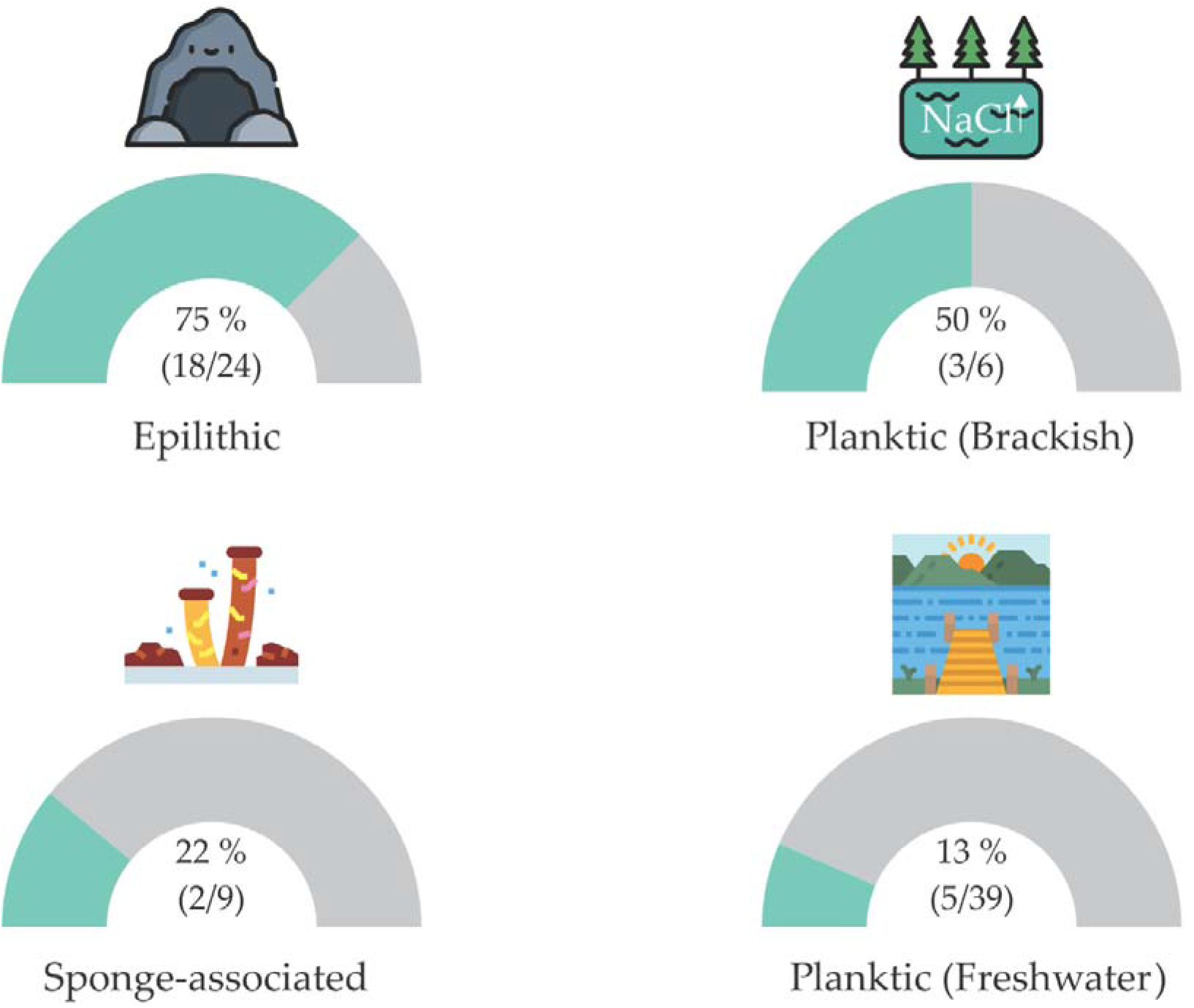
Cyanogenic relative frequency of cyanobacteria strains amongst different lifestyles presented with Gauge charts.

### 3.2 Polyphasic Taxonomy

According to the combined morphology (Figure 2) and the phylogeny based on 16S-23S ribosomal RNA (rRNA) and *cpcBA*-internal genetic spacer (IGS) regions (Figure 3), the strains, isolated in this study, positive for HCN production were classified into 18 genera and 20 taxa belonging to Chroococcales, Synechococcales, Oscillatoriales, Nostocales, and Pleurocapsales. Nine strains were identified to the genus level (*Allinostoc*, *Chloroglea*, *Oculatella*, *Komarekiella*, *Myxosarcina*, *Phormidium*, *Radiocystis*, and *Tolypothrix*), whereas the rest were identified up to the species level.

**Figure 2.**
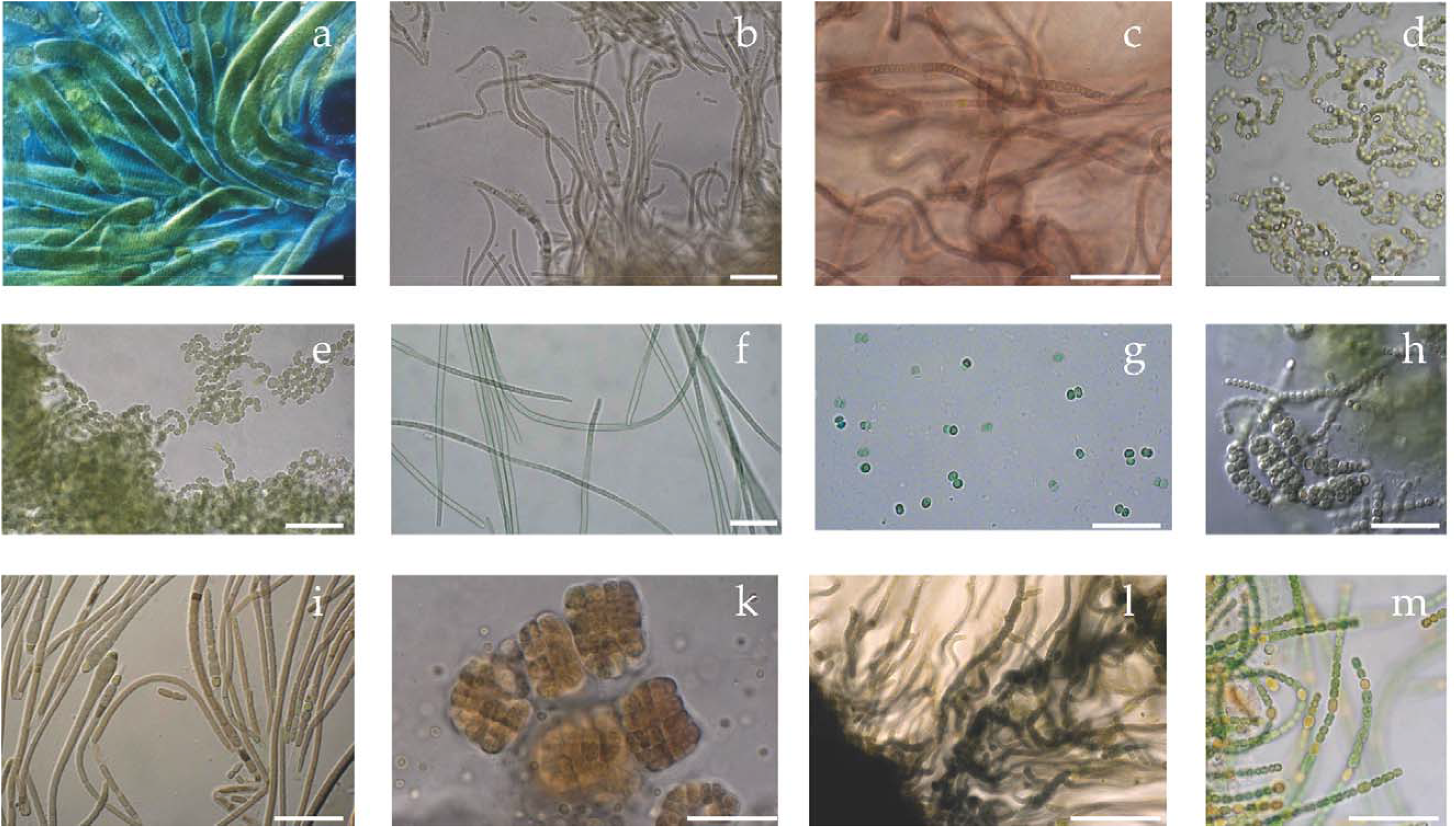
Microphotographs of strains, isolated in this study, representing 12 genera of cyanobacteria strains capable of producing HCN. (a) *Gloeotrichia echinulata* TAU-MAC 3718; (b) *Kovacikia muscicola* TAU-MAC 0518; (c) *Oculatella* sp. TAU-MAC 3318; (d) *Aliinostoc* sp. TAU-MAC 3117; (e) *Nostoc muscorum* TAU-MAC 1518; (f) *Desertifilum tharense* TAU-MAC 1517; (g) *Radiocystis* sp. TAU-MAC 1214; (h) *Komarekiella* sp. TAU-MAC 0117; (i) *Calothrix thermalis* TAU-MAC 1117; (k) *Myxosarcina* sp. TAU-MAC 3418; (l) *Geitleria calcarea* TAU-MAC 0118; (m) *Nodularia harveyana* TAU-MAC 0817. Scale bar = 10 μm.

**Figure 3:**
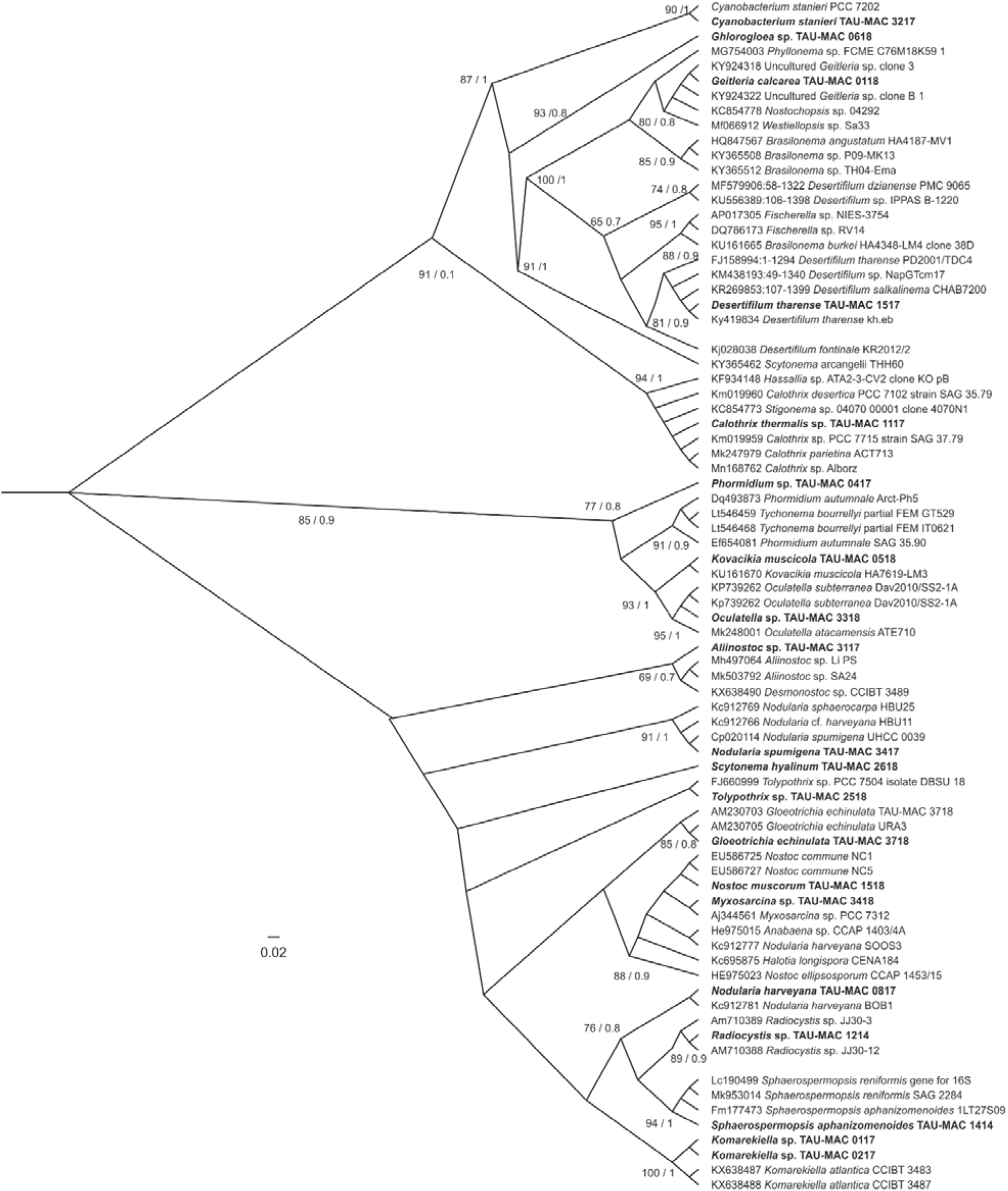
Phylogenetic tree based on 16S-23S rRNA and cpcBA-IGS sequences of HCN-producing cyanobacteria strains, isolated and described in this study. The phylogenetic tree was reconstructed using the Maximum-Likelihood (ML) and the Bayesian Inference (BI) analyses. ML topology is demonstrated. Numbers above branches indicate the bootstrap value (as percentages of 1,000 replications) for ML method and the posterior probabilities for BI method, respectively. Strains of the present study are indicated in bold. Bar represents 0.020 nucleotide substitutions per site.

Strains isolated from terrestrial caves formed clades with no more than two sequences, due to limited available sequences for cyanobacteria derived from these type of environments (Figure 3). Specifically, *Geitleria calcarea* TAU-MAC 0118 clustered with two uncultured *Geilteria* sp. sequences, whilst *Alliinostoc* sp. TAU-MAC 3117 was placed separately, outside two uncultured *Alliinostoc* sp. sequences. *Komarekiella* strains, isolated from a show cave in Chios Island in NE Aegean Sea, clustered together, separately of two *Komarekiella atlantica* strains, isolated from a tropical rainforest in Hawaii.

### 3.3 NifH Amplification

The *nifH* gene fragment was amplified (Figure S2) in 53 of the total 78 (67 %) cyanobacteria strains tested for HCN production (Figure S2). The *nifH* was present in all HCN-producing cyanobacteria strains, as well as in 25 more strains. All strains, classified to Nostocales order, carried the *nifH* gene fragment, whilst *nifH* was also amplified in non-heterocytous genera.

## 4. Discussion

### 4.1 Cyanogenic Cyanobacteria

In this study we demonstrate the production of cyanide from 28 cyanobacterial strains classified to 22 planktic and epilithic genera inhabiting freshwater, brackish, marine, terrestrial, and sponge-associated habitats. HCN production is stimulated mainly from histidine (Pistorius and Voss, 1977) and other aromatic amino acids is a general reaction catalysed by D-amino acid and L-amino acid oxidoreductases. The stoicheiometry of this process was investigated by Gewitz et al. (Gewitz et al., 1980) using snake venom L-amino acid oxidase and horseradish. Under optimal conditions, this system converted 72% of the added histidine into cyanide. Other products of the reaction were imidazole- Caldehyde, imidazole4carboxylic acid, CO, ammonia, water and imidazole acetic acid. The amount of CO, produced equalled the quantity of histidine oxidized and the sum of the ammonia plus cyanide formed. The cyanide production pathway is conserved in different type of organisms and it mainly contains an L-amino acid and its oxidase (Knowles and Bunch, 1986).

The production of cyanide by algae was first demonstrated by Gewitz et al. (Gewitz et al., 1976b) in 1976 in the green-alga *Chlorella vulgaris*. They showed that HCN is formed in small amounts when extracts were illuminated in the presence of O_2_ and supplemented with Mn^2+^ ions and peroxidase. In their study, a large number of amino acids were tested as possible precursors of HCN and D-Histidine was found to be the best promotor of cyanogenesis. Other aromatic amino acids could also promote cyanide formation (Gewitz et al., 1976a). These experiments also showed, that even though extracts of *Chlorella vulgaris* released HCN from amygdalin, a plant cyanogen, the HCN produced under oxidative conditions was not formed in this way (Gewitz et al., 1976b). Interestingly, the New Zealand spinach plant has a similar system for producing HCN (Gewitz et al., 1976a), as well as it was able to form cyanogenic glucosides in the grana. Further studies revealed details of the mechanism, used by the extracts of *Chlorella vulgaris* to convert histidine to HCN. Pistorius et al. (Pistorius and Voss, 1977) showed that a soluble protein, plus a component of the particulate fraction of extracts, were necessary for cyanogenesis. The soluble protein was found to be a D-amino acid oxidase, a flavoprotein, partially purified by Pistorius and Voss (Knowles and Bunch, 1986).

The production of cyanide in cyanobacteria has been reported only once in *Anacystis nidulans* (=*Asterocapsa nidulans*), *Plectonema borganum,* and *Nostoc muscorum* 40 years ago (Pistorius et al., 1979; Vennesland, 1981). Pistorius et al. (Pistorius et al., 1979) reported that histidine could stimulate HCN production. Larger quantities of HCN were produced if peroxidase, or certain redox metals, were also present suggesting that either the amino acid oxidase is located in the outer part of the cells, or the imino acid intermediate is excreted (Vennesland, 1981). Not surprisingly, the cells carried an L-amino acid oxidase (Pistorius and Voss, 1980). This enzyme has two subunits, each of 49,000 molecular weight, and contains one molecule of FAD per molecule of enzyme. It acts only on basic amino acids. Histidine is oxidized at a much slower rate. It is inhibited by divalent cations and orthophenanthroline. This latter observation implied a requirement for a metal ion, like zinc (Vennesland, 1981). In *A. nidulans* the L-amino acid oxidase has been reported to be associated with photosystem I (Pistorius and Voss, 1982).

All HCN-producing strains had the *nifH* gene, indicating that are capable of N_2_ fixation, and thus suggesting that their ability to fix nitrogen could be linked to the production and simultaneously reduction of HCN, although this needs to be confirmed. Nitrogen fixation has been linked with cyanide not only in living organisms (Knowles and Bunch, 1986), but also through synthetic methods for terminal nitride functionalization via conversion to the rare methoxymethyl imido unit (Curley et al., 2011). The ability of strains to fix nitrogen and produce cyanide seems to be not strictly related with heterocyte formation, as *nifH* gene fragment was also present in non-heterocytous cyanobacteria strains that produce nitrogen. Non-heterocytous cyanobacteria can fix nitrogen either in dark (Bergman et al., 2006) or in light combined with mechanisms for protecting the O_2_-labile nitrogenase (Berrendero et al., 2016). HCN can inhibit a wide range of metabolic processes (Knowles and Bunch, 1986), but the most pertinent effect in photosynthetic micro-organisms seems to be the inhibition of the reduced form of nitrate reductase (Lorimer et al., 1974).

An interesting relation was noticed concerning the relative frequency of HCN production amongst different cyanobacteria lifestyles. In the present study, cyanobacteria strains, isolated from epilithic mats, found to be more capable of HCN production compared to strains from aquatic ecosystems. Since cyanogenesis is a defence mechanism widely distributed in the plant kingdom and present in many major crop species (Thompson et al., 2016; Alberti et al., 2017), our results could possibly imply an unidentified chemical cue, released by different grazers, that triggers cyanobacteria to use HCN as a defence mechanism. Indeed, in environments where cyanobacteria are vulnerable to grazers such as rotifers or ciliates a rapid defence system should be favoured evolutionary (Wolfe, 2000). Even though no direct observations of activated chemical systems in unicellular organisms have been made, there are several examples of activated microbial defence reactions, which might serve as conceptual models for such systems (Mazard et al., 2016). One cannot exclude that physical contact with a grazer might stimulate cyanobacteria cells to produce compounds with a potential defensive role. Yang et al. (Yang and Kong, 2012) observed that the cyanobacterium *Microcystis aeruginosa* remaining under *Ochromona*s grazer pressure not only created colonies but also increased the amount of produced exopolysaccharides.

Cyanobacteria that thrive under extreme and diverse conditions tend to accumulate more compatible solubles such as sucrose, trehalose, glucosylglycerol, and glycine (Soule and Garcia-Pichel, 2019). However glycine accumulation could be toxic for the cyanobacterium and should be moderated (Eisenhut et al., 2007). Castric et al. (Castric, 1983)suggested that cyanogenesis is a response to a build-up of the intracellular glycine concentration. The key primary metabolic enzymes are serine hydroxymethyltransferase and glycine cleavage enzyme which catalyse conversion of serine into glycine and glycine into CO_2_ and ammonia, respectively. This could be the link between HCN production and cyanobacteria that pose a non planktic lifestyle.

### 4.2 Biodiversity

The polyphasic taxonomy applied to the strains of this study revealed taxa known to be part of bloom-forming communities (*Sphaerospermopsis* and *Nodularia),* rock-dwelling communities (*Scytonema* and *Tolypothrix*), and hot spring cyanobacteria mats (*Desertifilum* and *Gloeotrichia*) (Dadheech et al., 2014; Mazur-Marzec et al., 2015; Gkelis et al., 2017; Joanna and Andrzej, 2018). Our results revealed the presence of taxa not previously described from Greek habitats (Gkelis et al., 2016), such as *Allinostoc* (Saraf et al., 2018) and *Oculatella* (Osorio-Santos et al., 2014) and taxa previously described only from the tropical zone like *Komarekiella* and *Kovacikia* (Miscoe et al., 2016; Hentschke et al., 2017). Furthermore, the *Geitleria calcarea* strain isolated in this study, to the best of our knowledge, is the first isolate in world’s cyanobacteria culture collections depositories (Friedmann, 1979; Coute, 1989), whilst *Chlorogloea* sp. TAU-MAC 0618 consists the first bacteria isolate from an anchialine type environment.

The strains TAU-MAC 0817 and 3417 isolated from two brackish environments belong to species *Nodularia harveyana* and *Nodularia spumigena*, respectively. These strains constitute the first isolates of *Nodularia* strains in Greece, whilst in the Mediterranean there are only five strains of *Nodularia* isolated from Turkey (Akcaalan et al., 2009). The scarce records of *Nodularia* across Mediterranean could be linked with the absence of a high number of brackish environments. Anagnostidis (Anagnostidis, 1968),decades ago, reported the occurrence of *N. spumigena* and *N. harveyana* in Greece, (Gkelis et al., 2016). The largest research activity on the genus *Nodularia* occurs on the Baltic Sea, where *Nodularia spumigena* forms highly toxic blooms with significant effects on aquatic and non-aquatic organisms (Sivonen et al., 1989; Finni et al., 2001; Mazur-Marzec et al., 2015). Strain TAU-MAC 1517, isolated from a thermal site in Iceland was classified to the species *Desertifilum tharense* in both phylogenetic and morphological analysis, exhibiting the trichome’s “anchored” end, that discriminates it from *Microcoleus* and *Geitlerinema* (Dadheech et al., 2014). *Desertifilum tharense* has been recorded in India, Kenya, Mexico, Greece, Mongolia, and China (González-Resendiz et al., 2019). The presence of *Desertifilum tharense* in Iceland, a different ecological niche, supports the theory that despite its wide ecological span, the genus *Desertifilum* remains genetically stable (Sinetova et al., 2017; González-Resendiz et al., 2019).

Several strains of this study, such as *Geitleria calcarea*, *Kovacikia muscicola*, *Komarekiella* sp., *Scytonema hyalinum*, and *Tolypothrix* sp. were isolated from extensive dark-green coverings dominated by cyanobacteria like *Phormidium*, *Tolypothrix, Scytonema*, and *Geitleria* species. Cyanobacteria are considered the pioneering inhabitants in cave colonization (Joanna and Andrzej, 2018). They prevail in the cave entrances compared to the other microalgae (Mulec and Kosi, 2009) by colonizing various parts of the cave entrances, where biodiversity is the lowest (Vinogradova N. et al., 1998). Cyanobacteria represent the first photosynthetic colonizers on the calcareous surfaces usually thriving both as epiliths and as endoliths (Lamprinou et al., 2009). Lamprinou et al. (Lamprinou et al., 2013) observed predominance of Oscillatoriales group over Chroococcales in caves; our results show that also many Nostocales species form part of the cave communities, several of which, are understudied. For example, the *Komarekiella* sp. strains we isolated here are reported for the first time as cave inhabitants. Hentschke et al. (Hentschke et al., 2017),based on intensive examination of species life cycle, proposed the new genus *Komarekiella* and classified the strains from both Hawaiian and Brazilian rainforests in the single *Komarekiella atlantica* species. Our phylogenetic analysis of 16-23S and *cpcBA*-IGS sequences, as well as the different climatic zone and habitat, suggests that strains TAU-MAC 0117 and 0217, may belong to a new taxon, thus further research is required to describe it. Concerning *Geitleria calcarea* TAU-MAC 0118, as in the original description of the species, the most obvious finding is its apparent inability to produce heterocytes naturally (Coute, 1989). In our phylogenetic analysis, *Geitleria calcarea* TAU-MAC 0118 clustered with the only two available *Geitleria* sequences in GenBank, belonging to a non-cultured *Geitleria* species, derived from a population genetics study.

Terrestrial meteoric water mixed with saline groundwater resembles in a two-layered circulation, in estuaries termed as subterranean estuaries or anchialine (meaning near the sea) environments (Moore, 1999). *Chlorogloea* sp. TAU-MAC 0618 consists, to the best of our knowledge, the first bacterium isolate from this type of environments, as the research concerning bacteria, is focused mainly in community analysis, either by metagenomics (Brankovits et al., 2017), or fluorescence microbial profiling analysis (Seymour et al., 2007; Krstulovi et al., 2013).

## 5. Conclusions

The analysis of cyanobacteria strains from various environments revealed a high degree of biodiversity, deserving further research, whilst molecular data from strains may provide new information for cyanobacteria diversity, such as the isolation and characterisation of species like *Geitleria calcarea* and *Kovacikia muscicola* that provided a missing study material for underexplored cyanobacteria. We demonstrated the production of HCN by cyanobacteria spanning a wide taxonomic range across different habitats and lifestyles. The high percentage of epilithic cyanobacteria producing HCN suggest that it may be also used as defence mechanism, therefore exploiting cyanide production very differently compared to other cyanogenic microbes. All HCN-producing cyanobacteria carried the *nifH* gene fragment highlighting the complex mechanisms between nitrogen fixation and HCN production. The widespread cyanide production we report here calls for further research to investigate the significance cyanide metabolism has in the cycling of carbon and nitrogen, especially as plants, and probably cyanobacteria, both produce and catabolize cyanide.

## Supplementary Materials

Table S1: Complete list of TAU-MAC cyanobacteria strains used in this study, along with their description reference and the result of HCN Production. Plate position refers to Fig. S1 and strain number refers to Fig. S2. Strain number column refer to the number assigned to each strain for the PCR amplification of *nifH* gene fragment, Table S2: PCR primers used the phylogenetic analysis of cyanobacteria strains of TAU-MAC culture collection. Table S3: GenBank accession numbers for TAU-MAC strains used in the phylogenetic analysis **Figure S1**: Feigl-Anger Papers Of The HCN producing strains. Blue dot is indicating the production Of HCN. Plate position per strain refers to table S3. Con indicates positive control - *Trifollium repens*, Figure S2: PCR amplification of *nifH* gene fragment in the 78 TAU-MAC cyanobacteria strains tested for HCN production. Sample numbers refer to Table S3; + and − indicate positive and negative control, respectively; L indicates DNA ladder.

## Acknowledgments

The authors would like to thank Mrs. Kristel Panksep and Dr. Alejandro Martinez for providing sample material from Lake Peipsei (Estonia) and Túnel de la Atlántida (Lanzarote, Canary Islands), respectively. The authors would like also to thank Mr. Georgios Kakkos and Mr. Kostas Magkos for providing permission for sampling inside the Perama and Olympon Show Caves, respectively. Mr. Eythimios Parisis and Ms. Christina Skodra are acknowledged for providing help for the isolation of some strains as part of their final-year project supervised by S.G.

