## Supplemental Tables and Figures for "Cyano-assassins: Widespread cyanogenic production from cyanobacteria"

Article

**Table S1**: Complete list of TAU-MAC cyanobacteria strains used in this study, along with their description reference and the result of HCN Production. Plate position refers to Fig. S1 and strain number refers to Fig. S2.

| Strain Number | Strain | Reference | HCN-Production (Plate Position) |
| --- | --- | --- | --- |
| 1 | *Aliinostoc* sp. TAU-MAC 3117 | This study | Positive (A11-I) |
| 2 | *Anabaena* cf. *oscillarioides* TAU-MAC 0199 | (Gkelis et al., 2019) | Negative |
| 3 | *Anabaena* sp. TAU-MAC 1918 | This study | Negative |
| 4 | *Anabaenopsis elenkinii* TAU-MAC 0414 | This study | Negative |
| 69 | *Aphanothece* sp. TAU-MAC 4818 | This study | Negative |
| 27 | *Calothrix epiphytica* TAU-MAC 0399 | (Gkelis et al., 2019) | Negative |
| 28 | *Calothrix* sp. TAU-MAC 2318 | This study | Negative |
| 5 | *Calothrix thermalis* TAU-MAC 1117 | This study | Positive (C11-I) |
| 6 | *Chlorogloea* sp. TAU-MAC 0618 | This study | Positive (C8-I) |
| 29 | *Chlorogloeopsis fritschii* TAU-MAC 0599 | (Gkelis et al., 2019) | Negative |
| 72 | *Chroococcus* sp. TAU-MAC 1818 | This study | Negative |
| 7 | *Cyanobacterium stanieri* TAU-MAC 3217 | This study | Positive (D5-I) |
| 30 | *Cylindrospermopsis raciborskii* TAU-MAC 1414 | (Panou et al., 2018) | Negative |
| 8 | *Desertifilum tharense* TAU-MAC 1517 | This study | Positive (E4-I) |
| 53 | *Desmonostoc muscorum* TAU-MAC 0699 | (Gkelis et al., 2019) | Negative |
| 9 | *Geitleria calcarea* TAU-MAC 0618 | This study | Positive (A4-I) |
| 52 | *Geitlerinema* sp. TAU-MAC 0315 | This study | Negative |
| 13 | *Gloeocapsa* sp. TAU-MAC 1118 | This study | Negative |
| 10 | *Gloeotrichia echinulata* TAU-MAC 3718 | This study | Positive (G8-I) |
| 31 | *Hapalosiphon* sp. TAU-MAC 0115 | This study | Negative |
| 72 | *Jaaginema* sp. TAU-MAC 0110 | (Gkelis et al., 2019) | Positive (A3-II) |
| 78 | *Jaaginema* sp. TAU-MAC 0210 | (Gkelis et al., 2019) | Negative |
| 65 | *Jaaginema* sp. TAU-MAC 2210 | (Gkelis et al., 2019) | Negative |
| 11 | *Komarekiella* sp. TAU-MAC 0117 | This study | Positive (C4-I) |
| 12 | *Komarekiella* sp. TAU-MAC 0217 | This study | Positive (C6-I) |
| 14 | *Kovacikia muscicola* TAU-MAC 0518 | This study | Positive (H5-I) |
| 77 | *Leptolyngbya* sp. TAU-MAC 3218 | This study | Negative |
| 48 | *Leptothoe kymatousa* TAU-MAC 1215 | (Konstantinou et al., 2019) | Negative |
| 49 | *Leptothoe kymatousa* TAU-MAC 1615 | (Konstantinou et al., 2019) | Negative |
| 50 | *Leptothoe sithoniana* TAU-MAC 0915 | (Konstantinou et al., 2019) | Negative |
| 51 | *Leptothoe spongobia* TAU-MAC 1015 | (Konstantinou et al., 2019) | Negative |
| 15 | *Leptothoe spongobia* TAU-MAC 1115 | (Konstantinou et al., 2019) | Positive (C12-II) |
| 74 | *Limnothrix redekei* TAU-MAC 0310 | (Gkelis et al., 2019) | Negative |
| 75 | *Lyngbya* sp. TAU-MAC 4418 | This study | Negative |
| 76 | *Microcoleus* sp. TAU-MAC 2618 | This study | Negative |
| 59 | *Microcystis aeruginosa* TAU-MAC 0610 | (Gkelis et al., 2019) | Negative |
| 61 | *Microcystis* flos-aquae TAU-MAC 0410 | (Gkelis et al., 2019) | Negative |
| 63 | *Microcystis* flos-aquae TAU-MAC 1410 | (Gkelis et al., 2019) | Negative |
| 64 | *Microcystis* flos-aquae TAU-MAC 1510 | (Gkelis et al., 2019) | Negative |
| 66 | *Microcystis* flos-aquae TAU-MAC 1610 | (Gkelis et al., 2019) | Negative |
| 67 | *Microcystis* flos-aquae TAU-MAC 2010 | (Gkelis et al., 2019) | Negative |
| 68 | *Microcystis* sp. TAU-MAC 0710 | (Gkelis et al., 2019) | Negative |
| 70 | *Microcystis* sp. TAU-MAC 1710 | (Gkelis et al., 2019) | Negative |
| 71 | *Microcystis* sp. TAU-MAC 2110 | (Gkelis et al., 2019) | Negative |
| 73 | *Microcystis* sp. TAU-MAC 2310 | (Gkelis et al., 2019) | Negative |
| 56 | *Microcystis* sp. TAU-MAC 2410 | (Gkelis et al., 2019) | Negative |
| 55 | *Microcystis viridis* TAU-MAC 1810 | (Gkelis et al., 2019) | Negative |
| 16 | *Myxosarcina* sp. TAU-MAC 3418 | This study | Positive (E2-I) |
| 57 | *Nodosilinea* sp. TAU-MAC 0104 | (Gkelis et al., 2019) | Negative |
| 17 | *Nodularia harveyana* TAU-MAC 0817 | This study | Positive (F9-I) |
| 18 | *Nodularia spumigena* TAU-MAC 3417 | This study | Positive (F5-I) |
| 32 | *Nostoc calcicola* TAU-MAC 2918 | This study | Negative |
| 33 | *Nostoc elgonense* TAU-MAC 0299 | (Gkelis et al., 2019) | Negative |
| 19 | *Nostoc muscorum* TAU-MAC 1518 | This study | Positive (A2-I) |
| 34 | *Nostoc oryzae* TAU-MAC 2610 | (Gkelis et al., 2019) | Negative |
| 35 | *Nostoc oryzae* TAU-MAC 2710 | (Gkelis et al., 2019) | Negative |
| 36 | *Nostoc* sp. TAU-MAC 0799 | (Gkelis et al., 2019) | Negative |
| 39 | *Nostoc* sp. TAU-MAC 0899 | (Gkelis et al., 2019) | Negative |
| 20 | *Oculatella* sp*.* TAU-MAC 3318 | This study | Positive (E8-II) |
| 21 | *Phormidium* sp. TAU-MAC 0417 | This study | Positive (A6-II) |
| 45 | *Planktothrix agardhii* TAU-MAC 0514 | This study | Negative |
| 60 | *Pseudanabaena* cf. *perscicina* TAU-MAC 1415 | (Konstantinou et al., 2018) | Negative |
| 46 | *Pseudanabaena limnetica* TAU-MAC 0614 | This study | Negative |
| 22 | *Radiocystis* sp. TAU-MAC 1214 | This study | Positive (D9-II) |
| 40 | *Rivularia atra* TAU-MAC 3618 | This study | Negative |
| 41 | *Rivularia* sp. TAU-MAC 4718 | This study | Negative |
| 62 | *Schizotrichaceae* sp. TAU-MAC 1315 | (Konstantinou et al., 2018) | Negative |
| 23 | *Scytonema hyalinum* TAU-MAC 2618 | This study | Positive (D9-I) |
| 24 | *Sphaerospermopsis aphanizomenoides* TAU-MAC 1414 | This study | Positive (A12-II) |
| 37 | *Stenomitos* sp. TAU-MAC 4318 | This study | Negative |
| 38 | *Synechococcus* cf. *nidulans* TAU-MAC 3010 | (Gkelis et al., 2019) | Negative |
| 43 | *Synechococcus elongatus* TAU-MAC 3217 | This study | Negative |
| 44 | *Synechococcus* sp. TAU-MAC 0499 | (Gkelis et al., 2019) | Negative |
| 54 | *Synechococcus* sp. TAU-MAC 0715 | (Konstantinou et al., 2018) | Negative |
| 25 | *Synechococcus* sp. TAU-MAC 0815 | (Konstantinou et al., 2018) | Positive (A8-II) |
| 26 | *Tolypothrix* sp. TAU-MAC 2518 | This study | Positive (B9-II) |
| 42 | *Trichormus variabilis* TAU-MAC 1614 | This study | Negative |
| 47 | *Trichormus variabilis* TAU-MAC 2510 | (Gkelis et al., 2019) | Negative |
| 58 | *Xenococcus* sp. TAU-MAC 0615 | (Konstantinou et al., 2018) | Negative |

**Table S2:** PCR primers used the phylogenetic analysis of cyanobacteria strains of TAU-MAC culture collection.

| **Primer** | **Target-gene** | **Sequence (5^’^ –3^’^ )** | **Size (bp)** | **Reference** |
| --- | --- | --- | --- | --- |
| PCbF  PCaR_mod | *cpcBA*-IGS | GGCTGCTTGTTTACGCGACA CCAGTTCCACCAGCAATCAG | 720 | (Neilan et al., 1995) (Manen and Falquet, 2002) |
| Cya106F  23S30R | 16S-23S rRNA | CGGACGGGTGAGTAACGCGTGA CTTCGCCTCTGTGTGCCTAGGT | 1850 | (Nübel et al., 1997) (Lepère et al., 2000) |
| nifHf nifHr | *nifH* | CGTAGGTTGCGACCCTAAGGCTGA GCATACATCGCCATCATTTCACC | 300 | (Gugger et al., 2005) |

**Table S3:** GenBank accession numbers for TAU-MAC strains used in the phylogenetic analysis.

| **Strain** | **Accession Number (16S rRNA)** | **Accession Number (ITS region)** | **Accession Number (*cpcBA*-IGS)** |
| --- | --- | --- | --- |
| *Aliinostoc* sp. TAU-MAC 3117 | MN145677 | MN145697 | MN153483 |
| *Calothrix thermalis* TAU-MAC 1117 | MN145678 | MN145698 | MN153484 |
| *Chrologloea* sp. TAU-MAC 0618 | MN145679 | MN145699 | MN153485 |
| *Cyanobacterium stanieri* TAU-MAC 3217 | MN145680 | MN145700 | MN153486 |
| *Desertifilum tharense* TAU-MAC 1517 | MN145681 | MN145701 | MN153487 |
| *Geitleria calcarea* TAU-MAC 0618 | MN145682 | MN145702 | MN153488 |
| *Gloeotrichia echinulata* TAU-MAC 3718 | MN145683 | MN145703 | MN153489 |
| *Komarekiella* sp. TAU-MAC 0117 | MN145684 | MN145704 | MN153490 |
| *Komarekiella* sp. TAU-MAC 0217 | MN145685 | MN145705 | MN153491 |
| *Kovacikia muscicola TAU-MAC 0518* | MN145686 | MN145706 | MN153492 |
| *Myxosarcina* sp. *TAU-MAC 3418* | MN145687 | MN145707 | MN153493 |
| *Nodularia harveyana* TAU-MAC 0817 | MN145688 | MN145708 | MN153494 |
| *Nodularia spumigena* TAU-MAC 3417 | MN145689 | MN145709 | MN153495 |
| *Nostoc muscorum* TAU-MAC 1518 | MN145690 | MN145710 | MN153496 |
| *Oculatella* sp*.* TAU-MAC 3318 | MN145691 | MN145711 | MN153497 |
| *Phormidium* sp. TAU-MAC 0417 | MN145692 | MN145712 | MN153498 |
| *Radiocystis* sp. TAU-MAC 1214 | MN145693 | MN145713 | MN153499 |
| *Scytonema hyalinum* TAU-MAC 2618 | MN145694 | MN145714 | MN153500 |
| *Sphaerospermopsis aphanizomenoides* TAU-MAC 1414 | MN145695 | MN145715 | MN153501 |
| *Tolypothrix* sp. TAU-MAC 2518 | MN145696 | MN145716 | MN153502 |

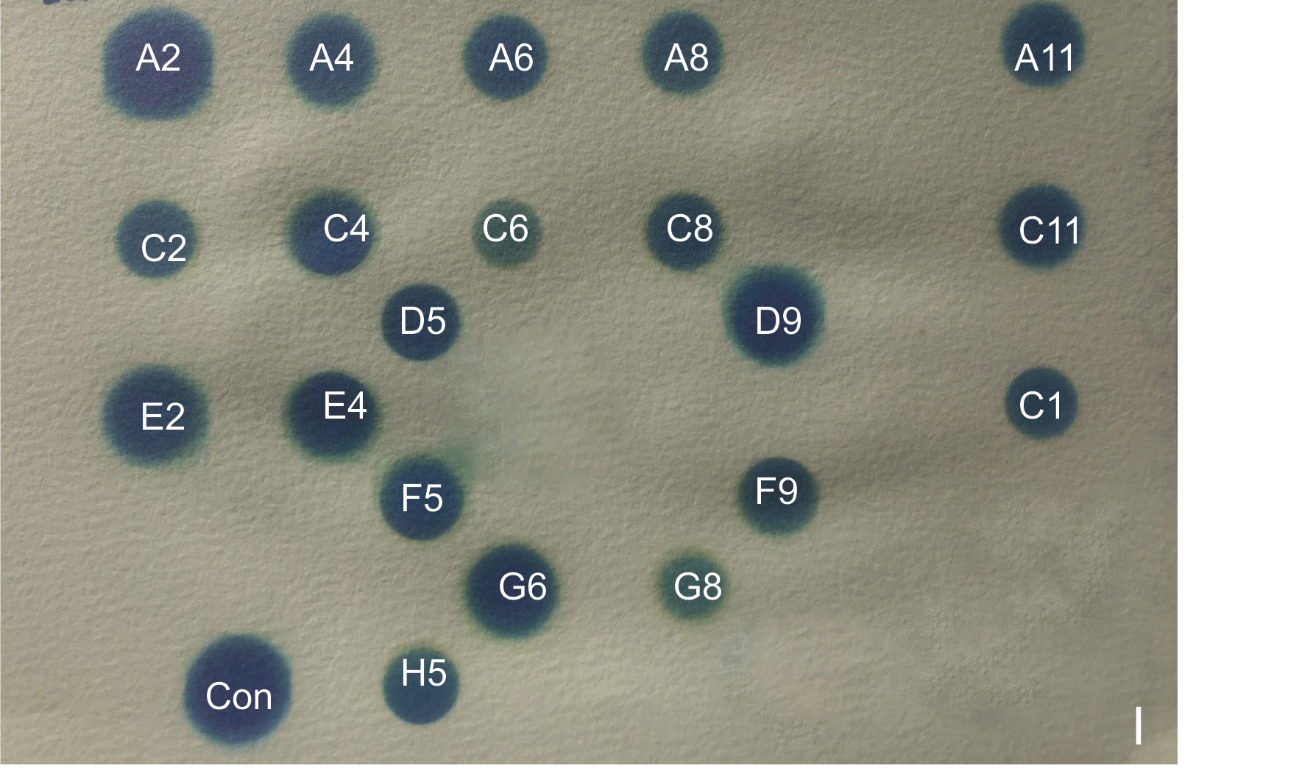

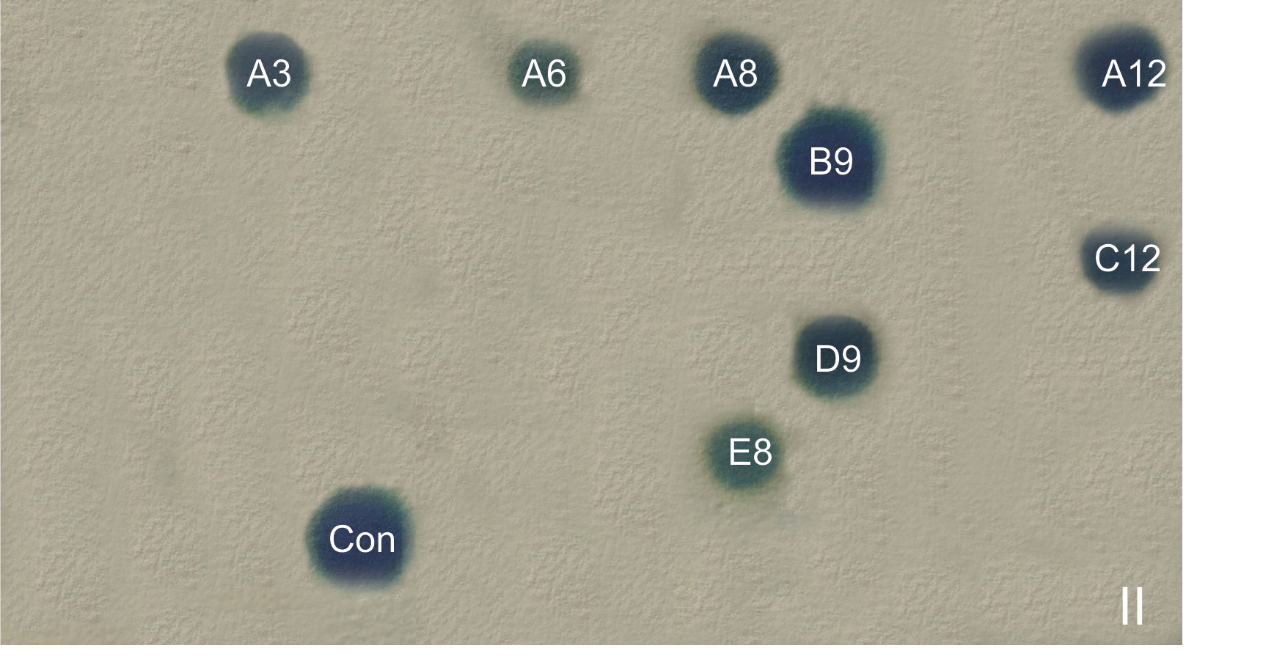

**Figure S1:** Feigl-Anger Papers of the HCN producing strains. Blue dot is indicating the production of HCN. Plate position per strain refers to table S3. Con indicates positive control - *Trifollium repens*.

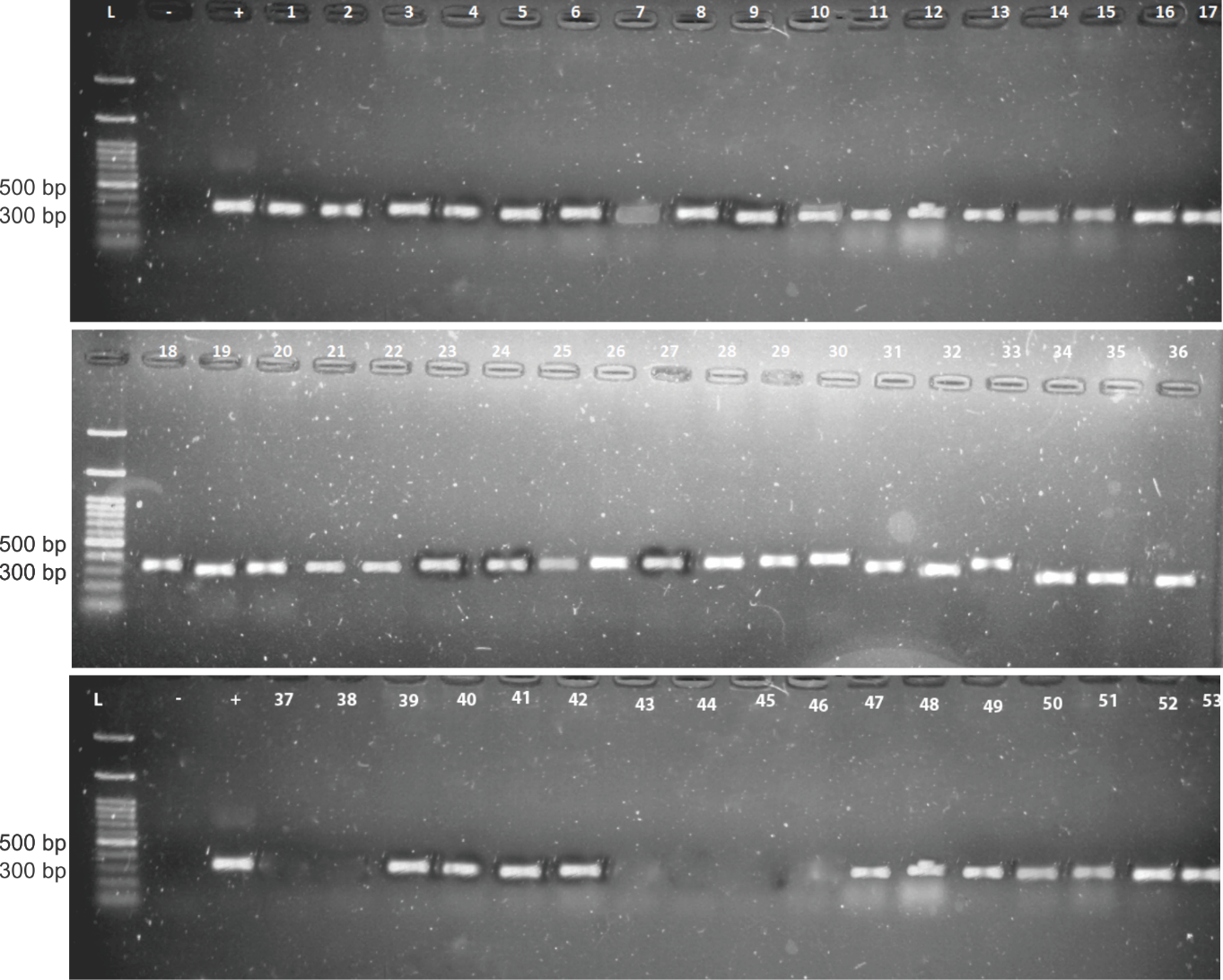

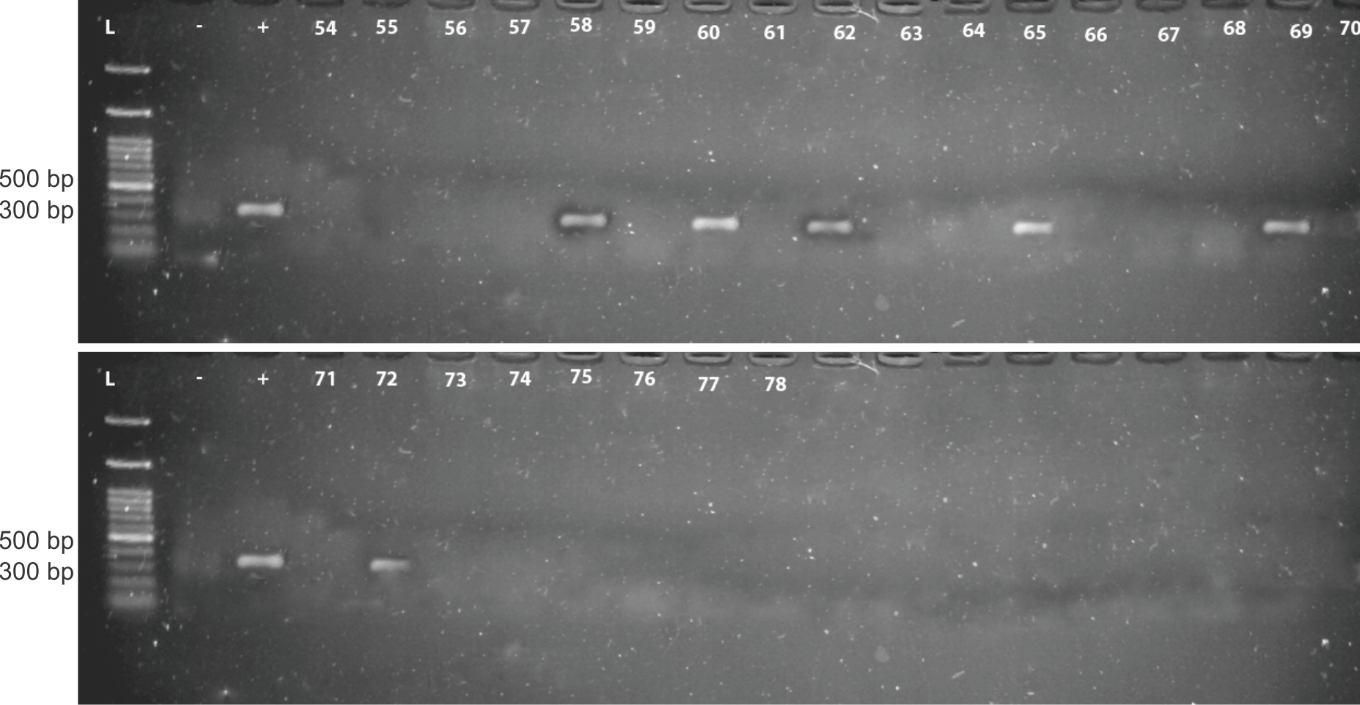

Figure S2: PCR amplification of *nifH* gene fragment in the 78 TAU-MAC cyanobacteria strains tested for HCN production. Sample numbers refer to Table S3; + and – indicate positive and negative control, respectively; L indicates DNA ladder.
